# The Effects of Statistical Multiplicity of Infection on Virus Quantification and Infectivity Assays

**DOI:** 10.1101/343723

**Authors:** Bhaven Mistry, Maria R. D’Orsogna, Tom Chou

## Abstract

Many biological assays are employed in virology to quantify parameters of interest. Two such classes of assays, virus quantification assays (VQA) and infectivity assays (IA), aim to estimate the number of viruses present in a solution, and the ability of a viral strain to successfully infect a host cell, respectively. VQAs operate at extremely dilute concentrations and results can be subject to stochastic variability in virus-cell interactions. At the other extreme, high viral particle concentrations are used in IAs, resulting in large numbers of viruses infecting each cell, enough for measurable change in total transcription activity. Furthermore, host cells can be infected at any concentration regime by multiple particles, resulting in a statistical multiplicity of infection (SMOI) and yielding potentially significant variability in the assay signal and parameter estimates. We develop probabilistic models for SMOI at low and high viral particle concentration limits and apply them to the plaque (VQA), endpoint dilution (VQA), and luciferase reporter (IA) assays. A web-based tool implementing our models and analysis is also developed and presented. We test our proposed new methods for inferring experimental parameters from data using numerical simulations and show improvement on existing procedures in all limits.

## I. INTRODUCTION

Understanding viral dynamics is an important task in medicine, epidemiology, public health, and, in particular, for the development of antiviral therapies and vaccines. Drugs that hinder viral infection include blockers of viral entry into the host cell [1–6] and inhibitors of genetic activity and protein assembly inside the cytoplasm and nucleus [7–9]. Mechanistic models of drug action have recently emerged as useful tools in helping design ad-hoc experiments to study drug efficacy and in interpreting results [10–13]. Mathematical models typically assume prior knowledge of given physical quantities pertaining to the virus, host cell, or the biological assay being studied. Once these parameters are assigned, viral and cell population dynamics and their statistical properties can be predicted. Among the different experimental assays, one often seeks to evaluate the number of virus particles in a stock solution or the number of viruses that have successfully infected host cells [6, 14**?** –18].

In the case of virus quantification assays (VQA), performing repeated controlled experiments on viral dynamics or comparing results across multiple studies requires knowing how many viruses are present in the initial stock solution of each experiment [4, 5]. Furthermore, antigens that induce immune responses against viral infections may be engineered from viral components such as capsid proteins, viral enzymes, and genetic vectors [19], and may be used in the development of vaccines. Being able to determine the exact number of virus-derived antigens helps control the efficacy of vaccines and optimize yield [20–22].

Given the central role of VQAs, several assays have been designed to estimate viral particle counts. These include plaque [23] and endpoint dilution [22, 24] assays, which will be discussed in more detail in the remainder of this work. For now, we note that these assays involve repeatedly diluting an initial solution of virus particles in the presence of a layer of plated cells, until viral concentrations are low enough that the dynamics of an individual virus can be extrapolated. At these low particle counts, however, the discrete nature of the infection process cannot be neglected and can cause substantial discrepancies when replicating experiments. Average quantities are not necessarily representative, and a more in-depth approach in quantifying virus-cell interactions is necessary.

Infectivity assays (IA), on the other hand, aim to quantify the number of viruses that have successfully infected host cells under varying antiviral drug environments [14–16]. IAs may measure the total transcription activity across all cells, such as the luciferase reporter assay [15, 25], or may count the number of host cells that were successfully infected, such as the enzyme-linked immunosorbent assay (ELISA) and the immunofluorence assay with fluorescence activated cell sorting (FACS) [4, 14, 15, 26]. These assays are performed using undiluted solutions with large numbers of viral particles, reducing stochastic variability. The average number of viruses that infect a cell is estimated as the ratio of the number of viruses in solution to the number of plated cells, a quantity known as the multiplicity of infection (MOI) [18]. However, each cell may be infected by different numbers of viruses distributed around the average given by the MOI. In these cases, one may be interested in the complete probability distribution for the number of virus infections in each plated cell and in the related statistical variance.

In this paper, we derive a probability model for the distribution of viral infections per host cell, which we call the statistical multiplicity of infection (SMOI). The SMOI can be used as a starting point to help estimate the number of viral particles in solution in VQAs, and to determine a viral strain’s ability to successfully infect host cells in IAs. In Section II A, we present the mathematical foundations for the SMOI in the two experimentally relevant parameter regimes of small and large viral particle counts and derive a probability model for the total number of infected cells under any dilution level. In Section IIIA, we apply our models to the plaque assay and formulate a new method of analyzing plaque count data. In Section IIIB we employ a special case of the derived probability distribution to the endpoint dilution assays and compare our results to those arising from traditional titration techniques such as the Reed and Muench [27] and SpearmanKarber methods [28]. In Section III C, we use the large particle limit of our model to describe the luciferase reporter assay. Lastly, a discussion of our results, a side-by-side comparison with existing methods, and a link to web-based data analysis tools are provided in Section IV. Mathematical appendices and further discussion of experimental attributes such as cell size variability, coinfection, viral interference, and optimal experimental design using parameter sensitivity analysis are presented in the Supplemental Information (SI).

## II. METHODS

### A. Probabilistic Models of Statistical Multiplicity of Infection (SMOI)

A typical viral assay is initiated by laying a monolayer of *M* cells on the bottom of a microtiter well, as illustrated in Fig. 1 [17, 23, 24]. Although variability exists among experiments, *M* is often set within the range of 10^4^–10^5^ [14, 25] and is assumed to be a known experimental parameter. A supernatant containing *N*_0_ virus particles in the range of 10^5^–10^7^ [23–25], is then added to the microtiter well. While, theoretically, all *N*_0_ particles are capable of infection, not all will successfully infect a cell. Since infection of a host cell requires a complex sequence of biochemical processes that may include receptor binding, membrane fusion, reverse transcription, nuclear pore transport, and DNA integration [10, 18], virus particles that fail at one or several of these sequential steps lead to abortive infections. To differentiate, the particles that do succeed are called infectious units (IU) or plaque forming units (PFU). We will denote the number of IUs as *N ≤ N*_0_. Depending on the strain of virus, the particular experimental protocol used, and specific conditions of the assay, the random quantity *N* is distributed according to *N*_0_ and the overall effective probability that an arbitrary viral particle successfully infects a host cell. A proxy that is typically used in place of this effective probability is the “particle to PFU ratio” *Q*, an experimentally determined parameter that quantifies, on average, the minimum number of particles required to ensure at least one infected cell [29, 30]. *Q* is often treated as an *a priori* measured quantity, primarily associated with the particular strain of virus being studied. Low values of *Q*, such as with poliovirus (*Q* = 30) [30], have a high likelihood of successful infection compared to viruses with large *Q*, such as HIV-1 (*Q* = 10^7^) [31]. Thus, the reciprocal *Q*^−1^ can be interpreted as the probability for a single virus to infect a host cell. Assuming an initial stock of *N*_0_ particles, the discrete probability density function of *N* is

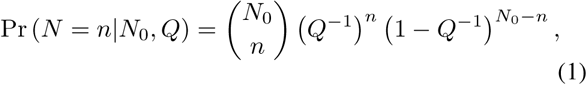

which defines a binomial distribution with parameters *N*_0_ and *Q*^−1^. Although we assume *Q* to be *a priori* known, in actuality, the probability of a virus successfully infecting a host is highly dependent on the methods used to harvest the virus stock, the experimental parameters of the assay, the host receptor concentrations and binding rates, and the dynamics of the physiological processes leading to infection [29, 32]. A thorough investigation into these processes would be necessary to mechanistically model *Q* and is outside of the scope of this paper. However, we will discuss in Section IV how, with direct measurements of certain other parameters, especially *N*_0_, our derived methods may also be used to infer *Q*.

**FIG. 1:**
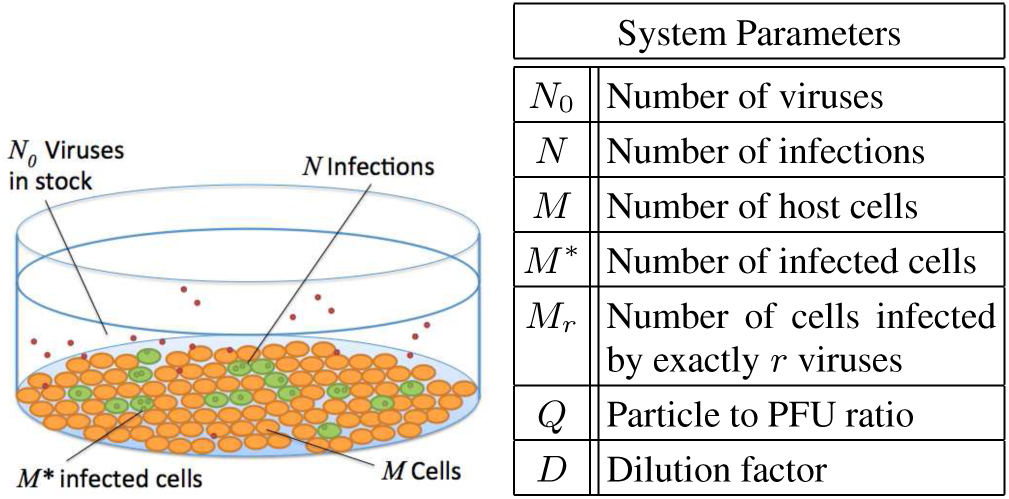
A typical assay includes a plate of *M* host cells inoculated with a solution of *N*_0_ viruses. Each viral particle has some probability of infection and the total number *N* of infections are distributed to the *M*\* infected cells. The probability of infection is roughly estimated with the reciprocal of the *a priori* measured particle to PFU ratio *Q*.

We assume each viral particle in solution acts independently of others and that host cell infection attempts are random events. At high ratios *N*_0_*/M* of particles to cells, a quantity referred to as the “multiplicity of infection” (MOI), it becomes increasingly probable for more than one IU to infect the same host cell. We define *M*_0_ as the count of cells not infected by any IU, *M*_1_ as the count of cells infected by exactly one IU, up to *M_N_*, the number of cells infected by all *N* IUs. The statistical multiplicity of infection (SMOI) is defined as the ensemble of cell counts 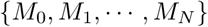. Note that two constraints must hold: 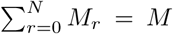 to account for all infected and un-infected cells, and 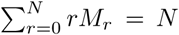 for conservation of the total number of IUs. If we assume all *M* cells are of identical size and volume, they carry equal probability of being infected by a particular virus. Thus, evaluating the probability distribution that *M_r_* takes on the value *m_r_* reduces to the well-known occupancy problem of randomly placing balls into identical urns [33] and we derive

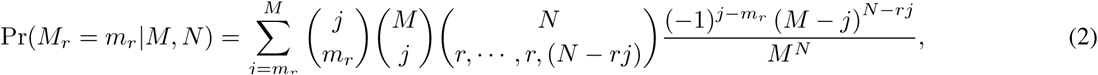

where the *r* term is repeated *j* times in the lower argument of the multinomial coefficient. The derivation of Eq. 2 is detailed in Appendix A in the SI and an investigation into the effects of inhomogeneous cell sizes is presented in Appendix B. Furthermore, in Appendix A, we derive the expected value and variance of *M_r_* as

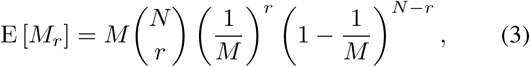

and

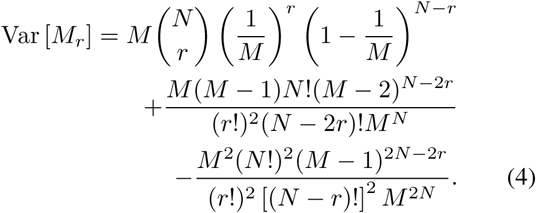

The first and second multinomial expressions enumerate the degeneracy of how the *M* identical cells are distributed across the configuration 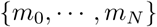 and how the *N* identical IUs are chosen for those cells respectively. Although the second expression in Eq. 5 is more succinct, it must be explicitly conditioned on the constraints 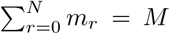 and 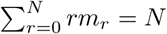.

The expressions in Eqs. 2 and 5 provide an exact discrete description of the stochasticity of the MOI, but are computationally expensive to evaluate for large values of *N* and *M*. In a typical virology experiment, the number of viral particles *N*_0_ and host cells *M* are large enough for certain asymptotic methods to be applicable. Furthermore, for intermediate values of *Q*, and based on Eq. 1, the expected number of IUs *N* would be similarly large. We can thus take the mathematical limit *N*, *M* → *∞* while keeping the ratio 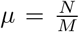 fixed and approximate Eq. 2 as:

Note that the variance is equal to the expected value with two additional correction terms that cancel each other as *N* and *M* increase, indicating the probability distribution of *M_r_* is Poisson-like for large *N* and *M*. A plot of a representative probability distribution and a test of agreement between our analytical result and numerical simulation is provided in Fig. 2.

**FIG. 2:**
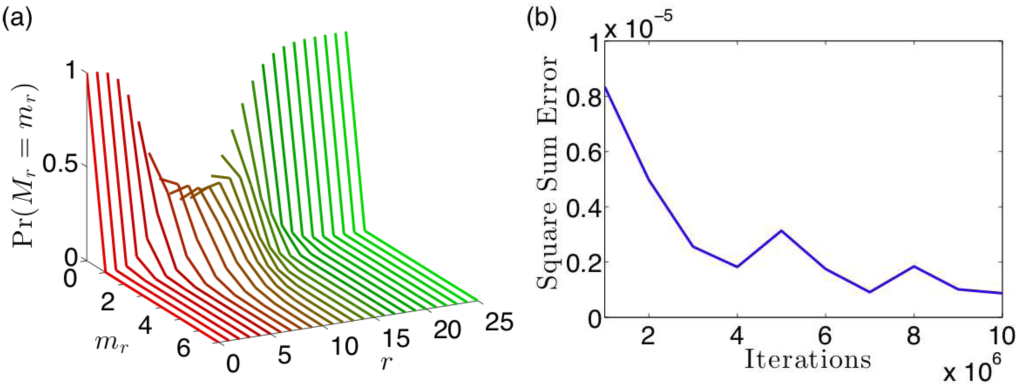
(a) A collection of curves of the probability of finding *m*, cells that have been infected by exactly *r* IUs given a total number of IUs *N* = 100 and a total number of cells *M* = 10 using Eq. 2. With *N/M* = 10, we expect very few cells to be uninfected, resulting in the probability distribution concentrated close to 0 for low values of *r*. Similarly, we expect few cells to be infected by a very large number of IUs, accumulating the probability distribution close to 0 for large *r*. Only at intermediate values of *r* ≈ *N/M* = 10 we observe a Poisson-like distribution. (b) We perform a numerical study to show empirically that our analytical result in Eq. 2 matches the statistical frequency of virus-cell counts from a simulation of *N* = 100 IUs being randomly assigned to *M* = 10 cells. The square sum error between the simulated proportions and the analytical result was calculated with increasing numbers of iterations of the simulation. For iterations around 10^6^, our square sum error is on the order of 10^−6^, indicating strong agreement between our model and simulation.

We also derive the joint probability 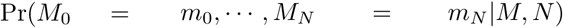 that the SMOI 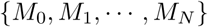 takes on the set of values 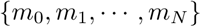 as

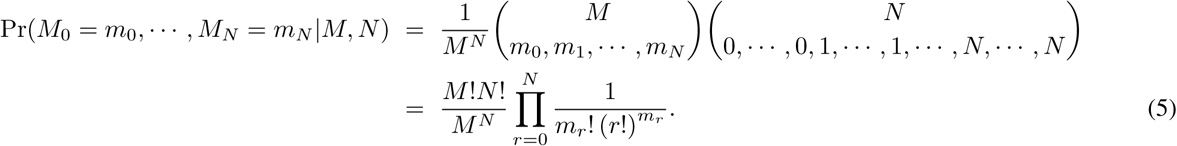

approximate Eq. 2 as:

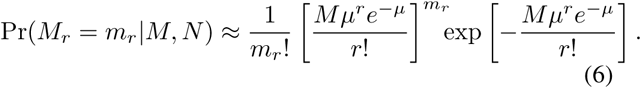

Eq. 6 implies that *M_r_* is Poisson-distributed with mean and variance

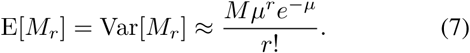

A mathematical justification of Eq. 6 is given in Appendix A and comparisons of Eq. 6 and the analytical result in Eq. 2 to simulations are shown in Fig. 3. Under the same large *M*, *N* limit and using Eq. 6, we show in Appendix A

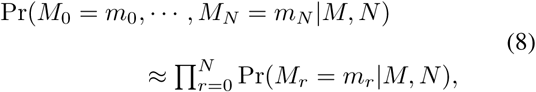

which implies that as *M*, *N* → *∞*, the random variables 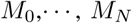 are independently distributed. In the next section, we will apply results of our probability model of SMOI to the case of a repeatedly diluted solution of virus particles, a procedure used in many VQAs.

**FIG. 3:**
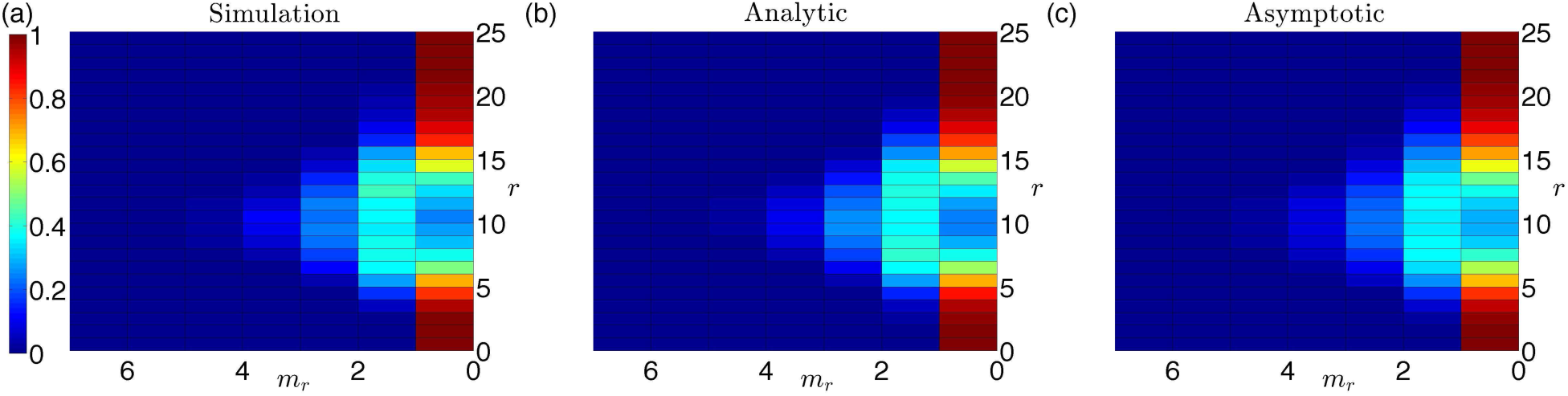
Heat maps of the probability distribution Pr (*M_r_* = *m_r_* | *M, N*) of finding *m_r_* cells that have been infected by exactly *r* IUs given a total number of viruses *N* = 100 and *M* = 10 cells. (a) The statistical frequency of virus-cell counts after simulating IUs randomly distributing to the *M* cells, averaged over 1000 iterations. (b) The analytical result obtained from Eq. 2. (c) The asymptotic approximation with *M* = 10 and **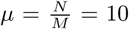**, using the expression in Eq. 6. There is close agreement between the simulated and analytical results. The relatively low values of *M* and *N* makes the asymptotic formula in Eq. 6 inappropriate for this parameter regime, explaining the discrepancy between the asymptotic result and the exact analytical result. However, it is noteworthy how qualitatively small that deviation is, which will continue to vanish as *M* and *N* increase in value.

### B. Serial Dilution

Low viral particle concentrations in assays are typically obtained via serial dilution processes in order to increase the sensitivity to individual viral infections [4, 23, 24]. The initial viral stock containing *N*_0_ particles is diluted by a fixed factor of *D* and the process is repeated *d*_max_ times. At each dilution number *d*, an assay can be performed to determine if the concentration of virus particles in the diluted solution is sufficient to generate a qualitative signal of infection, known as a “cytopathic effect” (CE). For example, the diluted stock can be administered *in vivo* to a model organism such as a mouse. The mouse’s death would indicate that at least one lethal unit of the virus was present at that dilution level. Alternatively, an *in vitro* assay can be carried out to measure a signal that, for example, quantifies the exact number of plated cells that were successfully infected. To model these assays, we first define *M*\* as the number of host cells infected by at least one IU and that are capable of producing new viruses. In Appendix A we derive the discrete probability density function for finding *M\** = *m* infected cells at a given dilution number *d* and find

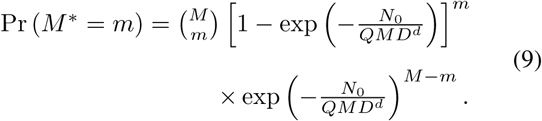

Eq. 9 shows that the number of infected cells *M*\* is binomially distributed with expected value

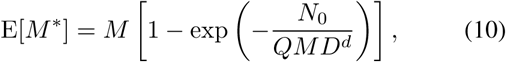

and variance

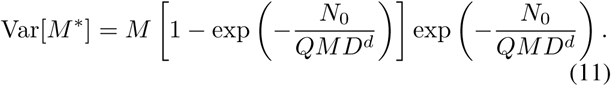

We can define the probability of observing a CE at dilution number *d* as the probability of finding one or more infected cells:

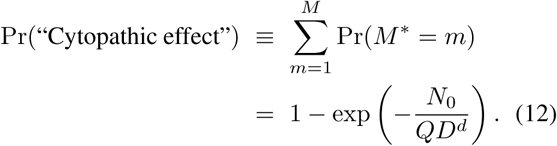

The definition we use in Eq. 12 assumes an *in vitro* assay that can exhibit a cytopathic signal after a single cell infection or more. For *in vivo* assays, the probability that *m* infected cells are sufficient for a CE will depend on many complex physiological factors such as immune pressure, in-host viral evolution, and virion burst size [34]. A plot of how the initial particle count *N*_0_ and dilution factor *D* effect the characteristic functional form of Eq. 12 are shown in Fig. 4. Although both Eqs. 9 and 12 assume each IU contains all viral genes required for in-host replication, an extended probability model that factors in genetic mutation and degradation is provided in Appendix C. Furthermore, for the case of retroviruses, infectious processes inside the host cytoplasm may be suppressed by previous infections, known as viral interference, and is explored in Appendix D. In Section III A, we will use Eq. 9 to analyze the plaque assay. Eq. 12 will be used for “binary” assays that are only concerned with the presence or absence of a CE such as the endpoint dilution assay, which we will explore in Section IIIB.

**FIG. 4:**
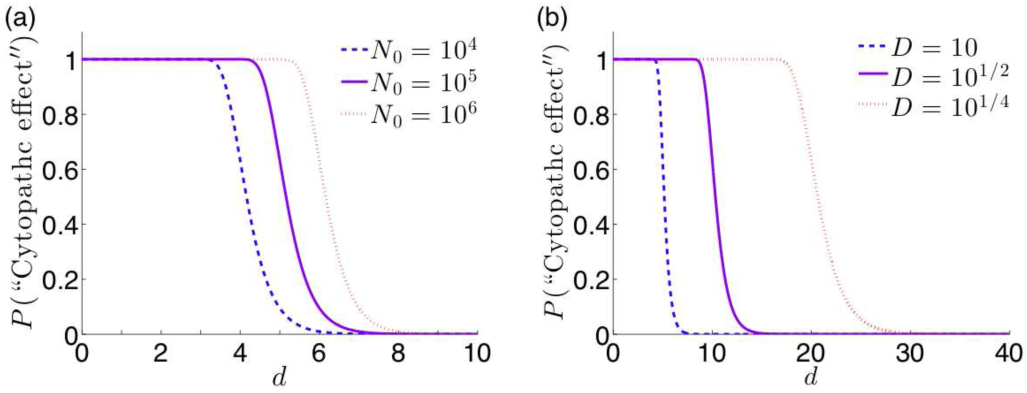
The probability of observing a cytopathic effect (CE) given in Eq. 12 as a function of the dilution number *d* and with *Q* = 1. (a) For *D* = 10, as the initial particle count *N*_0_ increases, the critical dilution moves toward higher *d*. (b) Common dilution factors include logarithmic dilution (*D* = 10), half-logarithmic dilution (*D* = 10^1/2^), and quarter-logarithmic dilution (*D* = 10^1/4^). Logarithmic dilution requires a lower number of dilutions to cause the characteristic decrease in probability, requiring less individual assays to perform. Quarter-logarithmic dilution, though requiring more dilutions, has a slower transition from high to low probability across *d*, making the assay less sensitive to experimental error or noise. The plot above can be used to quantify the tradeoffs between the choices of *D*.

## III. RESULTS AND DISCUSSION

### A. Plaque Assay

The plaque assay is an example of a virus quantification assay (VQA) where the objective is to infer the total number of viruses *N*_0_ present in a solution assuming the PFU to particle ratio *Q* has been independently measured and estimated [23, 24, 35]. After *d* serial dilutions, the viral stock is added to a monolayer of *M* cells and a layer of agar gel is added to the well to inhibit the diffusion of virus particles in the plate. If a virus successfully infects a host cell, the agar will limit the range of new infections to the most adjacent cells. Viral infection thus spreads out radially from the initial nucleation infection and forms a visible discoloration in the plate called a “plaque.” For high particle concentrations, the number of plaques formed may be large enough to cover the entire plate surface. After a sufficient critical dilution number *d_c_* however, the number of plaques formed are low enough to be visibly distinct and countable. For each dilution number *d*, the assay can be performed for *T* number of trials. The ‘signal’ data arising from the plaque assay *P_d,t_* is defined as the number of visible plaques counted, where 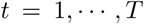 is the trial number. The standard method of obtaining an estimate 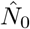 of the true particle count *N*_0_ is to apply the sample mean of the data 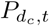 at the critical dilution level *d_c_* to the formula

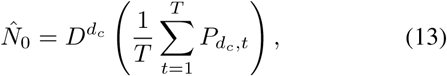

which posits that the average number of plaques is directly proportional to the particle count *N*_0_. Eq. 13 assumes that each infected cell corresponds to one IU, which is not necessarily true in the context of SMOI. Furthermore, although data corresponding to dilution numbers *d* < *d_c_* are unusable, data for *d* > *d_c_* corresponding to countable plaques are not used at all in Eq. 13.

In order to improve on Eq. 13 by using the entire set of plaque counts *P_d_*_,*t*_ for our estimate of *N*_0_, we propose a maximum likelihood estimation (MLE) scheme. Using the mathematical models derived above, we can construct an expression 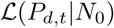 of the probability that the observed data *P_d_*_,*t*_ can be generated assuming a particular value for *N*_0_, known as a likelihood function. A value for *N*_0_ that maximizes 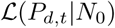 corresponds to the most probable estimate 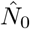 that could have generated the data. As each nucleation of a plaque corresponds to a distinct infected cell (and assuming that overlapping lesions of necrotic cells are still discernible as distinct plaques), we can equate *P_d_*_,*t*_ to the total number of successfully infected cells *M*\*. We will ignore the dynamics of coinfection and viral interference. Using Eq. 9, we propose the following likelihood function of the data given *N*_0_:

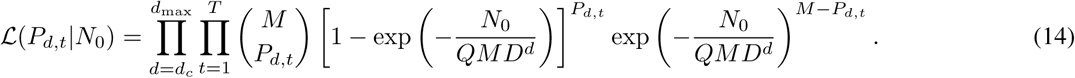

To obtan the MLE 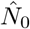, we take the derivative of the natural log of Eq. 14 with respect to *N*_0_ and set the result to zero to obtain

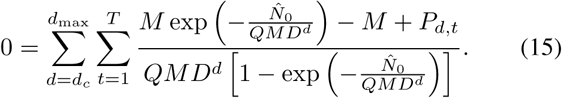

We can solve Eq. 15 for 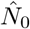 using numerical methods such as Newton-Raphson [36], an iterative scheme that approaches the solution of an equation asymptotically starting from an initial guess 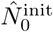. To increase the stability of convergence to the solution, we choose 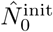 by equating the sample average of plaque counts 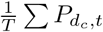 with the expected number of infected cells E[*M\**] in Eq. 10 at the critical dilution *d_c_* to derive

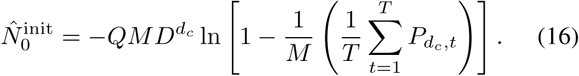

An example of raw plaque count data and the resulting estimates for *N*_0_ are given in Fig. 5. In order to quantify the relative improvement of the MLE of *N*_0_ over the standard method in Eq. 13, we simulate plaque assay data assuming a fixed, known *N*_0_ value. In our simulation, we use the models established in Section II A to sample the *N*_0_ particles according to Eq. (S9) in Appendix A to account for serial dilution and sample again the resulting particles according to Eq. 1 to obtain the number of IUs *N*. The IUs are distributed randomly to the *M* cells with equal probability and the resulting number of infected cells *M\** is recorded. Since plates of cells with too many infections render the number of plaques uncountable, a “countable plaque threshold” renders the data unusable when the number of infected cells exceed the threshold. Thus, the resulting plaque data *P_d_*_,*t*_ for a given dilution *d* and trial *t* is assigned the number of simulated infected cells if the latter is less than the given threshold. A scatter plot of the data *P_d_*_,*t*_ of one such simulation is shown in Fig. 6a and the corresponding likelihood function from Eq. 14 is plotted in Fig. 6b. Because the MLE method utilizes a full probabilistic model of the plaque count distribution instead of relying only on the expected value at the single critical dilution *d_c_*, it produces an estimate consistently closer to the original *N*_0_ that generated the data. To better quantify this property, in Appendix E we derive an asymptotic approximation of the variance of 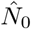 as

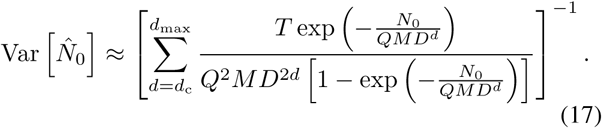

**FIG. 5:**
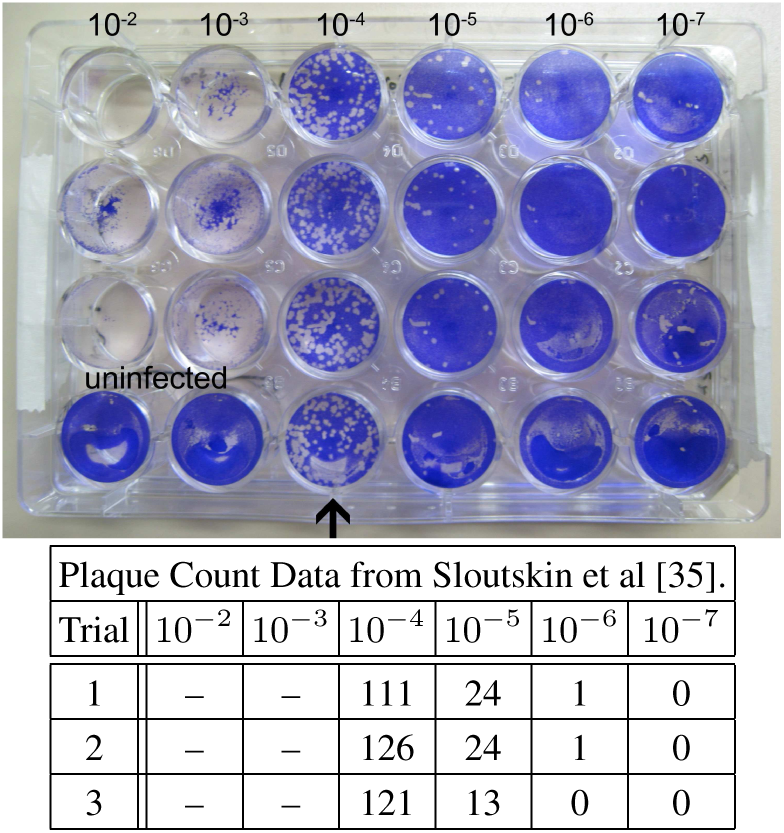
An example of raw plaque count data taken from Sloutskin et al. [35]. A viral solution was assayed in a plate of *M* = 3 × 10^5^ cells at dilution numbers *d* = 2, 3, 4, 5, 6, and 7 at a dilution factor of *D* = 10. The particle to PFU ratio is assumed to be *Q* = 1. For *T* = 3 separate trials, the number of plaques were counted at each dilution level. The bottom row of plates used as a control is ignored. For dilution numbers *d* = 2 and 3, the entire plate of cells show cytotoxicity so that the numbers of plaques were undiscernable and, thus, the countable data starts at *d_c_* = 4. For the old method featured in Eq. 13, the estimate for *N*_0_ is 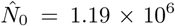 and for the MLE derived from Eq. 15, 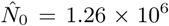. This results in a relative difference of 5.5%. Furthermore, when applying these parameters and 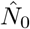 estimate to Eq. 17, we observe a 10.7% decrease in the estimate variation using the MLE technique.

**FIG. 6:**
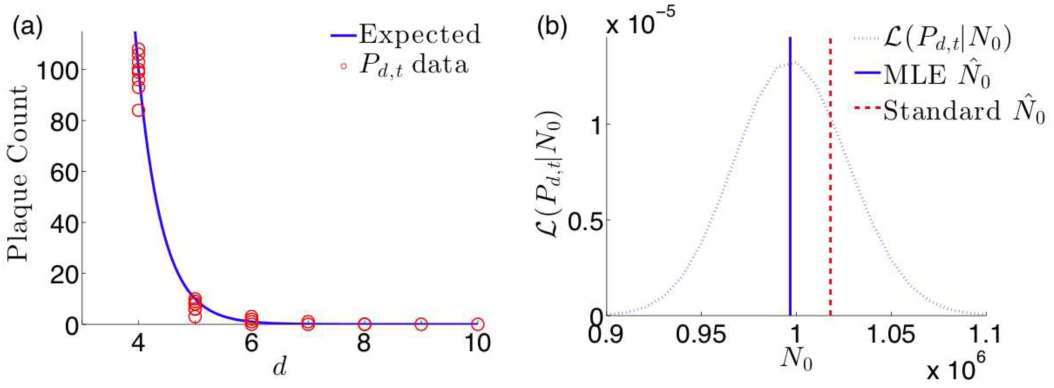
Results of plaque assay simulation for parameters *N*_0_ = 10^6^, *M* = 10^5^, *Q* = 1, *D* = 10, *d*_max_ = 10, and *T* = 10. (a) The scatter plot of simulated data *P_d_*_,*t*_ (circles) and the expected value of plaque counts as given by Eq. 10 show close agreement. (b) The likelihood function 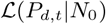 with respect to *N*_0_ using the same simulated data. The MLE obtained by iteratively solving Eq. 15 is 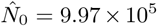 and is relatively closer to the true value of *N*_0_ than the estimate calculated from the standard method in Eq. 13 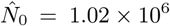.

The variance is an explicit function of *Q*, which is assumed to be *a priori* known. If there is uncertainty in the value of *Q*, Eq. 17 can quantify how sensitive the distribution of 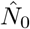 is to variation in *Q*, as shown in Fig. 7a. We can see that for small assumed *Q*, such as in poliovirus [30], error in this measurement can cause a large relative change in the accuracy of 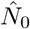. This type of sensitivity analysis on estimation variance can be done with any experimental parameter included in the likelihood function in Eq. 14. Furthermore, for directly controllable parameters, such as the serial dilution factor *D*, Eq. 17 can provide insight into optimizing the assay protocol, as shown in Fig. 7b. Although it is evident that small *D* would increase the accuracy of the 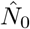 estimate, doing so requires more serial dilutions which increases the time and expense of the assay. Thus, our sensitivity analysis provides a quantitative method for making experimental design choices between minimizing uncertainty versus the cost of an assay protocol. Lastly, if we compute the variance of the standard method in Eq. 13 due to the known variance in the data *P_d_*_,*t*_, and compare with Eq. 17, we find, when using realistic parameter values from Fig. 5, the standard method results in a 10.7% higher variance than that of our method. Although the significance of the relative increase in precision of estimating *N*_0_ found using our method is highly dependent on the context of the experimental study for which the assay was performed, similar sensitivity analysis can be used to determine such tolerances.

**FIG. 7:**
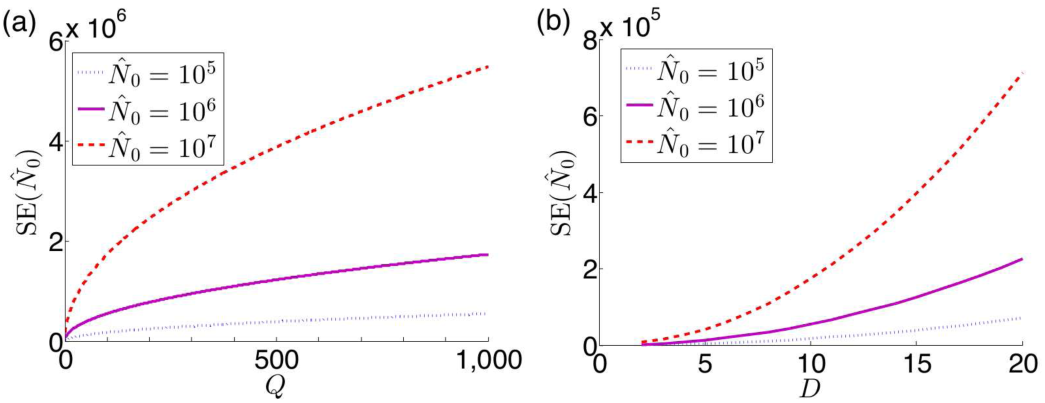
Approximations of the standard deviation 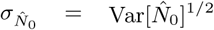 of maximum likelihood estimates for the plaque assay using Eq. 17 and parameters 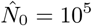, 10^6^, and 10^7^, *M* = 3 × 10^5^, *d*_c_ = 4, *d*_max_ = 7, and *T* = 3, corresponding to the assay displayed in Figure 5. (a) For *D* = 10, the standard deviation increases proportional to the square root of *Q*. (b) For *Q* = 1, we can see a low dilution factor *D* will increase the accuracy of the estimate 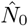.

### B. Endpoint Dilution Assay

Another widely used assay for quantifying the initial viral particle count *N*_0_ is the endpoint dilution or endpoint titration assay [22, 24, 37]. It is often used in place of the plaque assay as it can be more rapidly performed and is useful for viral strains that are unable to form plaques. Here, serial dilutions at a factor of *D* are employed and at every dilution number *d*, an assay is performed *T* times to test for a successful CE. The number *E_d_* of observed CEs among the *T* trials at a given dilution number *d* is recorded as the signal. For low dilution, we expect many cells to be infected and the probability of observing a CE, as shown in Eq. 12, is close to 1. If every trial of the assay is likely to display a CE, then *E_d_* is expected to be close to *T*. However, at high dilution, the probability in Eq. 12 rapidly decreases to 0, as shown in Fig. 4, and *E_d_* will be similarly small. For a large initial stock of viral particles *N*_0_, a larger dilution number *d* is needed to ensure the dramatic change in probability in Eq. 12. Thus, the critical dilution at which *E_d_* most rapidly decreases from *T* can be used to estimate the particle count *N*_0_. This occurs at the point of inflection when *d* = log*D* (*N*_0_*Q^−^*^1^) and corresponds to when the expected number of successful trials E[*E_d_*] = *T* (1 − *e*^−1^), as shown in Fig. 8.

**FIG. 8:**
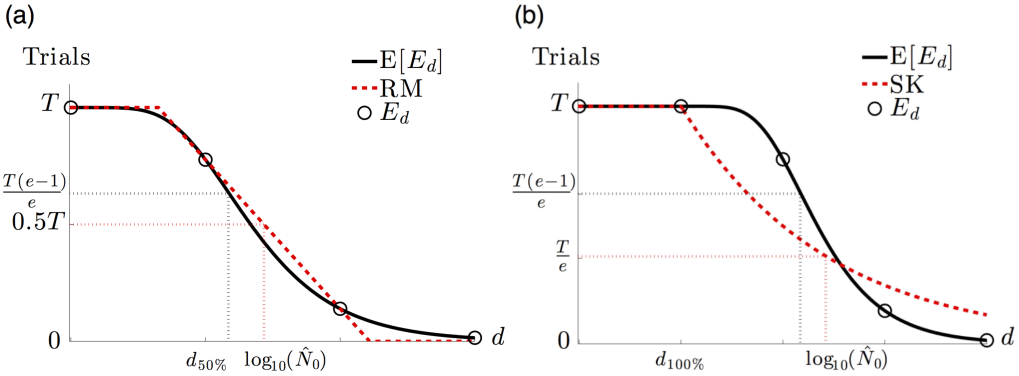
An illustration of the consistent overestimation of the Reed and Muench (RM) and Spearman-Karber (SK) methods using the expected curve E [*E_d_*] of CEs given *T* trials as a function of the dilution number *d* derived from Eq. 12. (a) The RM method approximates the steepest decent of the expectation curve with a line connecting the two data points 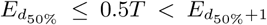. Because of the relative convexity of the expected curve, the linear approximation consistently rests above the curve and results in an overestimate of 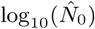. (b) From the last dilution *d*_100%_ such that all trials exhibit a CE, the SK method assumes an exponential decay of the expectation. Obtaining the characteristic decay rate of the exponential involves calculating the area under the curve, which is done numerically using the data *E_d_*. However, according to our model, many of the expected values of *E_d_* exist above the exponential, causing the numerical integration to overestimate the area and, thus, decay too slowly. This gradual decrease in the exponential curve results in a larger estimate of 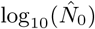.

One commonly used way to estimate *N*_0_ is the Reed and Muench (RM) method that utilizes the two dilution numbers that capture the greatest change in the data *E_d_* [27]. We first define a critical dilution number *d*_50%_ to be the largest dilution such that at least 50% of the trials exhibit a CE. The estimate 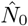 for the particle count *N*_0_ is given by

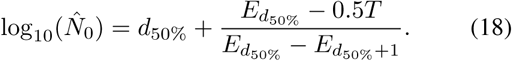

The RM method effectively attempts to approximate the steepest descent of the CE probability given in Eq. 12 with a line connecting the assay data at dilutions *d*_50%_ and *d*_50%+1_, as displayed in Fig. 8a. Unfortunately, this line always rests above the actual expectation curve of *E_d_*, so any estimate 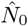 obtained from this method will overestimate the true *N*_0_. Another commonly used estimation scheme is the SpearmanKarber (SK) method which uses the critical dilution number *d*_100%_, the largest dilution such that 100% of trials exhibit a cytopathic effect [28, 37]. The SK estimate 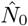 is given by

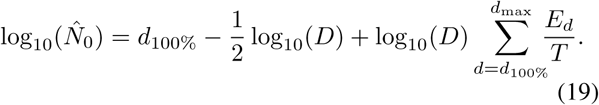

In this method, the downward slope for the expectation of *E_d_* is assumed to follow a decaying exponential starting at dilution *d*_100%_, as shown in Fig. 8b. The intention is to find the dilution at which *Te*^−1^ CEs are expected by calculating the area under the exponential curve, given by the summation term in Eq. 19. However, the actual values of *E_d_* will follow the expected curve from our model, leading to an overestimate of the area and, by extension, a larger value for 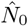. Both standard methods were derived from the heuristic observation that *E_d_* exhibits sigmoidal behavior as a function of the dilution number *d*, but an underlying probabilistic model was missing, resulting in consistent overestimation of the true *N*_0_. Furthermore, neither method uses the “particle to PFU ratio” *Q*, accounts for the stochasticity of serial diluting viral samples, considers the dynamics of SMOI, or employs the entire set of data *E_d_*.

We present an alternative way to infer *N*_0_ using Eq. 12 to establish a maximum likelihood estimation scheme. We restrict ourselves to *in vitro* assays in which a single infected cell is sufficient to display a CE. Then each cytopathic count is binomially distributed with parameters *T* and the probability given in Eq. 12. Thus, for a set of data 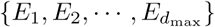, we propose the likelihood function

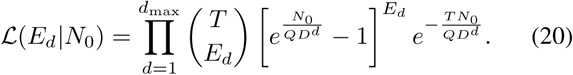

Eq. 20 is an expression of the probability of the data 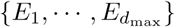 given the current assumed value of *N*_0_. To obtain the best estimate 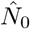 of *N*_0_, we maximize the likelihood function by taking the log and derivative of 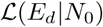 with respect to *N*_0_ and set it equal to zero to obtain

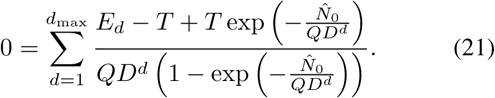

As with Eq. 15, solving Eq. 21 for 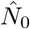 requires a numerical method such as Newton-Raphson. As an appropriate initial estimate for 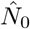, the formula

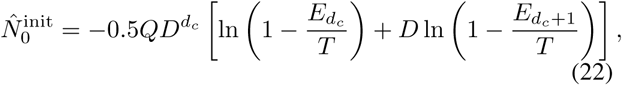

can be used, where *d_c_* is the largest dilution number such that at least half of the trials exhibit a cytopathic effect. Eq. 22 is the average of the *N*_0_ estimates at dilutions *d_c_* and *d_c_*_+1_ when setting the CE probability in Eq. 12 to 1*/*2. For a comparison of our MLE method with the RM and SK methods, we simulate data similar to that described in Section IIIA. Here we take the number of trials such that the simulated count of infected cells is greater than zero as the values of *E_d_* for a given dilution number *d*. We plot the likelihood from Eq. 20 and compare the MLE of *N*_0_ with those derived by the RM and SK methods in Fig. 9a. While both RM and SK estimate very similar values of 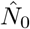, they both consistently over-estimate the *a priori* set *N*_0_ relative to the MLE method. This demonstrates the advantage of a probabilistic model for parameter inference over heuristically determined formulas.

**FIG. 9:**
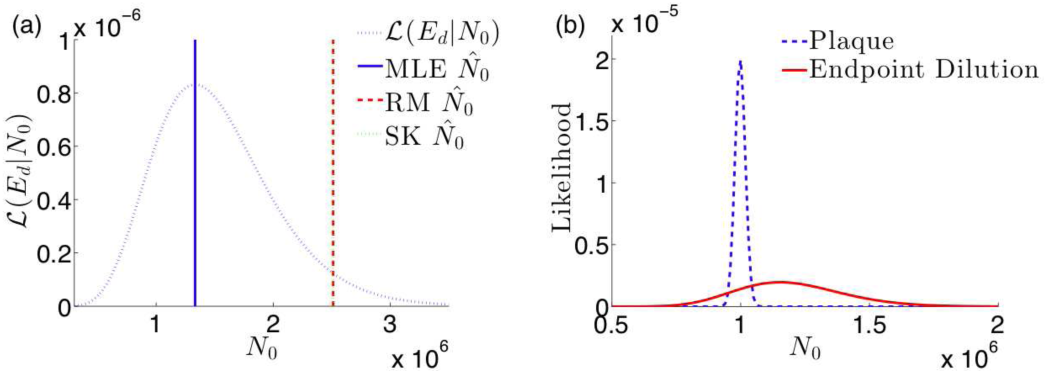
(a) The likelihood function 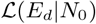 in Eq. 20 for the endpoint dilution assay and the corresponding maximum likelihood, Reed and Muench, and Spearman-Karber estimates given simulated data generated with *N*_0_ = 10^6^, *Q* = 1, *D* = 10, and *d*_max_ = 10. The estimates for maximum likelihood 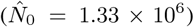, RM 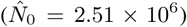, and SK 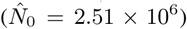 all overestimate *N*_0_, but the smaller relative error of the MLE is an improvement on the errors of the existing two methods. (b) The likelihood functions 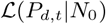 and 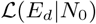 for the plaque and endpoint dilution assays respectively given simulated data. The data was generated with parameters *N*_0_ = 10^6^, *M* = 10^5^, *Q* = 1, *D* = 10^1/4^, *d*_max_ = 30, and a “countable plaque threshold” of 150. The plaque assay likelihood is concentrated close to the true *N*_0_ value while the endpoint dilution likelihood is far more spread out and overestimates *N*_0_. This direct quantitative comparison can inform an experimentalist when choosing between the two methods.

The expressions we derived in Eqs. 14 and 20 applied to simulated data can also help quantify tradeoffs in experimental design. As discussed above, there exist viruses that cannot form plaques, restricting the options of VQAs to endpoint dilution. However, for many cases, the choice between using one assay over the other can be one of convenience. More specifically, endpoint dilution assays can often be performed more rapidly than plaque assays. Using the same simulated data for both assays, we plot Eqs. 14 and 20 together in Fig. 9b. The plots clearly show the superiority of the plaque assay for estimating the viral stock number *N*_0_ in respect to both how close the MLE infers the true *N*_0_ value and the amount of variance in that estimate. While the amount of variability and error that is tolerable for an experiment may be context-dependent, the plots in Fig. 9b provide a quantitative way to differentiate between the two methods.

### C. Luciferase Reporter Assay

The luciferase reporter assay is commonly used to measure the infectivity of a viral strain. Here the ratio *µ* = *N/M* of total infections over the number of plated cells is estimated by measuring the transcription activity of viral proteins [14–16]. The reporter employs an oxidative enzyme luciferase that facilitates a reaction when introduced to the substrate luciferin, resulting in bioluminescence. The protocol begins with attaching the luciferase gene to the viral genome. The altered viral strain is cloned to a total particle count *N*_0_ which, in this case, is assumed to be fixed and known. The solution of viruses is added to a plated monolayer of *M* host cells. An incubation time is allowed for transcription of viral proteins and, incidentally, the luciferase enzyme. Subsequently all cells are lysed to release all cytoplasmic contents into the solution upon which luciferin is added. The oxidation of luciferin is facilitated by the luciferase enzyme and the resulting bioluminescence yields a measurable signal [38]. The light intensity is thus a measure of total transcription activity of the viral genome in all infected cells and can be used as a proxy for the total number of viruses *N* that successfully infected host cells.

Although there is stochasticity in transcription factor binding and, in the case of retroviruses, the number of integration sites on the host DNA, we will assume that each successful virus infection contributes one viral genome to be transcribed and each transcription occurs at a constant rate proportional to the total number of integrated viral genomes. Note that the limited number of transcription factors, ribosomes, and other cell machinery necessary to produce viral proteins and the luciferase reporter causes the production rate to saturate as the number of infecting viruses *r* per cell increases. Thus, transcription activity saturates with increasing number of infections *r*. We can model this effect by defining a monotonically increasing function *f* (*r*) representing the number of transcribed viral proteins when a cell is infected by *r* viruses over the course of the assay. Thus, for a given SMOI 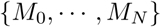, we will model the intensity signal *L* of the total luciferase reporter luminescence with

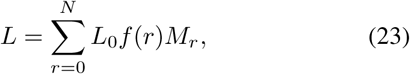

where *L*_0_ is the fluorescence intensity arising from a single luciferase reporter present in the solution. Although *f*(*r*) may take on many functional forms, a commonly used model for transcription factor kinetics is the Hill function [39] given by

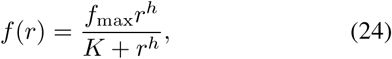

where *f*_max_ is the maximum transcription activity of luciferase, *h* is the Hill coefficient that effectively describes cooperative binding of multiple transcription factors at a promoter region, and *K* is an effective dissociation constant relating the binding and unbinding rates of transcription factor. The functional form of Eq. 24 accounts for the limited transcription machinery available for the multiple copies of viral genome present in the cell. In Fig. 10a we calculate the discrete probability distribution Pr(*L* = *ℓ*) by considering the cumulative weight of every allowable configuration of *N* viruses infecting *M* cells through Eq. 23.

**FIG. 10:**
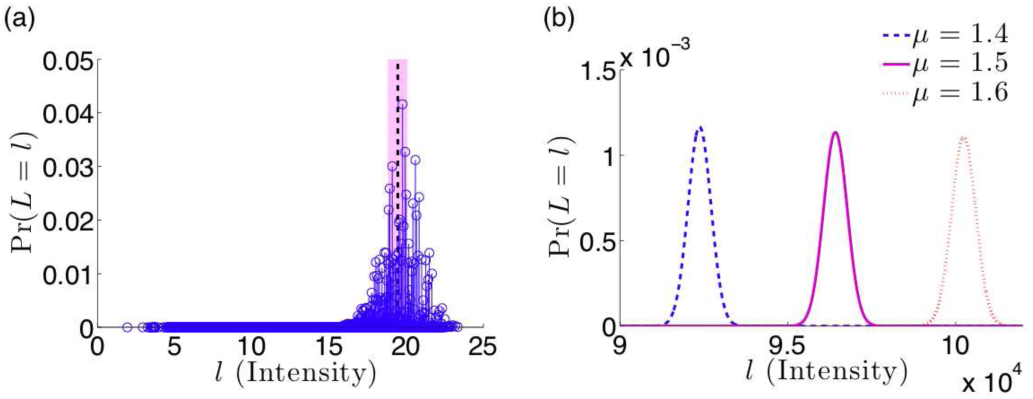
Probability distributions of the luciferase assay fluorescence intensity *L* from Eq. 23. (a) A toy example of a discrete probability distribution of allowable fluorescence intensities for *N* = 30 viruses infecting *M* = 20 cells. Due to the *M^N^* finite number of allowable configurations of the SMOI, there are a corresponding finite number of intensities with specific probabilities determined by Eq. 5 and represented by a unique circle. The parameters used for the reporter kinetics are *f*_max_ = 2, *h* = 1, *K* = 1 and *L*_0_ = 1. The mean intensity of the fluorescence signal is E[*L*] = 19.5, represented by the vertical dotted line, and variance Var[*L*] = 1.49, represented by the shaded region. (b) The normally distributed approximation of fluorescence intensity using *M* = 10^5^, *f*_max_ = 2, *h* = 1, *K* = 1 and *L*_0_ = 1. The distributions are plotted for *µ* = 1.4, 1.5, and 1.6 by computing the expected values E[*L*] = 9.23 × 10^4^, 9.64 × 10^4^, and 10^5^ and the variances Var[*L*] = 1.7 × 10^5^, 1.24 × 10^5^, and 1.3 × 10^5^ respectively.

Since luciferase reporter assays typically involve large values of initial virus count *N*_0_ and cell count *M*, we can use the asymptotic approximations in Eqs. 6 and 7 along with the Central Limit Theorem [40] to assume *L* is normally distributed with expected value

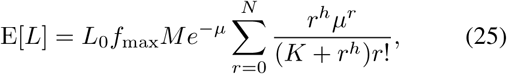

and variance

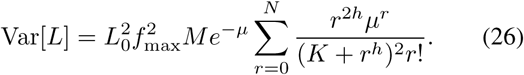

A visualization of the normal approximation of the probability distribution of *L* is shown in Fig. 10b. Furthermore, with Eqs. 25 and 26, we can derive the likelihood function 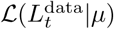 do the data 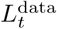, given *µ*

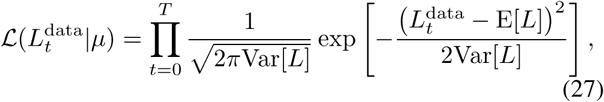

where 1 ≤ *t* ≤ *T* is the trial number. Due to the complicated functional form of the mean and variance of *L*, creating a maximum likelihood scheme to estimate *µ* from experimental data is intractable, so we use Eq. 25 by replacing the expected value with the experimental average of measurements 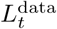. If we assume no cooperative transcription binding (*h* = 1), we solve for the estimate 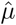 by applying the Newton-Raphson iterative method to the equation

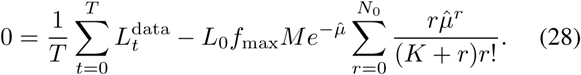

The typical method, under the assumption that luminescent intensity is proportional to the number of IUs *N*, is to use the sample mean via the formula 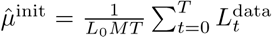. This standard approach fails to account for the effects of SMOI, but can be used to generate an initial guess for solving Eq. 28 iteratively. In order to compare the two estimates, we simulate data similar the descriptions in the previous two sections. Here, we do not dilute the initial particle count and, after distributing the *N* IUs to the *M* cells with equal probability, we compile the SMOI configuration and calculate 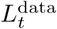 using Eq. 23. The results are shown in Fig. 11. The iterative method produces an estimate 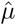 far closer to the true value of *µ* than the former method. A similar approach can be used to compare methods for alternative functional forms of the viral protein transcription dynamics described in Eq. 24.

**FIG. 11:**
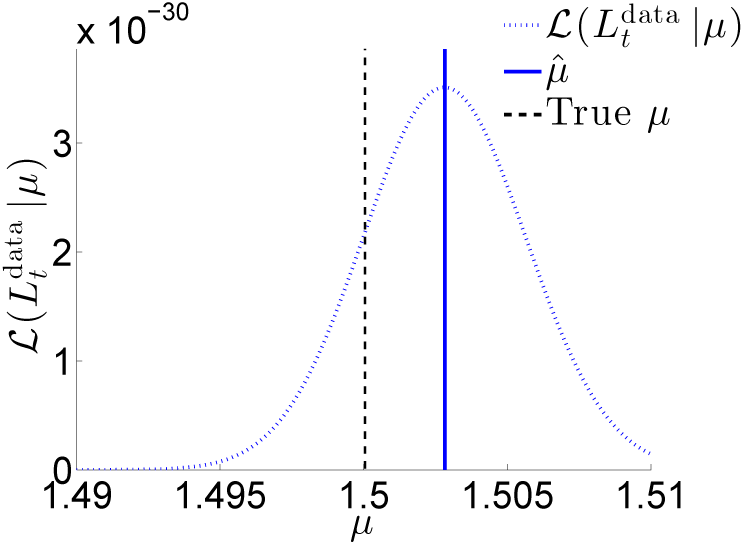
The likelihood function 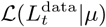 using Eq. 27 and simulated data. We set *µ* = 1.5 and assign other parameters with *M* = 10^5^, *f*_max_ = 2, *h* = 1, *K* = 1 and *L*_0_ = 1. The estimate derived from solving Eq. 28 is 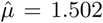 while the standard method based on the sample mean yields 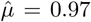, far lower than what is displayed in the plot.

### IV. CONCLUSION

In this work, we derived probability models that quantify the viral infectivity of host cells in an *in vitro* environment. By factoring in the stochastic nature of virus-host engagement, defective and/or abortive events, and the possibility of multiple infections of a single host, we defined the statistical multiplicity of infection (SMOI) and determined related probabilistic models. We analyzed two limiting regimes: small numbers of infecting viruses *N* and large *N*. For the low *N* regime, Eqs. 2 and 5 model how the limited number of infectious units are distributed amongst the *M* host cells. Alternatively, for large *N*, we showed the cell counts of the SMOI become statistically independent, as displayed in Eq. 8, and that they display a Poisson distribution (Eq. 6). Lastly, we explored the effects of serial dilution on the total number of infected cells and the probability of observing an infectious signal in Eq. 9.

Using our probability models along with reasonable assumptions of applied combinatorics and nonlinear inference, we analytically derived expressions for several virus assays to improve on existing methods of experimental data analysis. For virus quantification assays, serial dilution results in low numbers of viral particles. Using the appropriate probability model, we created new methods of estimating the particle count *N*_0_ in the initial viral stock for the plaque assay and the endpoint dilution assay. For measuring infectivity of a viral strain, the objective is to determine the effective multiplicity of infection *µ* = *N/M* as the ratio of successfully infecting viruses *N* and the total number of cells *M* included in the assay. As these assays operate under no dilution, we employed the large *N* limit probability model to analytically derive expressions for the luciferase reporter assay to estimate *µ*. A summary of each estimation method along with the most commonly used counterpart is displayed in Table I.

**TABLE I:** A summary of the analytically derived expressions used to analyze experimental results. For virus quantification assays, such as the plaque and endpoint dilution assays, one typically wishes to estimate the number of initial viral particles *N*_0_. For luciferase reporter infectivity assay, the ratio *μ* = *N/M* is desired. Our improved parameter estimation methods are listed next to standard methods currently used.

| Comparison of Virological Assay Analyses |  |  |
| --- | --- | --- |
| Assay (Parameter) | Standard Method | New Method |
| Plaque ($N_0$) | $\hat{N}_0 = D^{d_c} \left( \frac{1}{T} \sum_{t=1}^T P_{d_c,t} \right)$ | $0 = \sum_{d=d_c}^{d_{\max}} \sum_{t=1}^T \frac{M \exp\left(-\frac{\hat{N}_0}{QMD^d}\right) - M + P_{d,t}}{QMD^d \left[1 - \exp\left(-\frac{\hat{N}_0}{QMD^d}\right)\right]}$<br>Initial guess:<br>$\hat{N}_0^{\text{init}} = -QMD^d \ln \left(1 - \frac{1}{MT} \sum_{t=1}^T P_{d_c,t}\right)$ |
| Endpoint Dilution ($N_0$) | Reed and Muench:<br>$\log_{10}(\hat{N}_0) = d_{50\%} + \frac{E_{d_{50\%}} - 0.5T}{E_{d_{50\%}} - E_{d_{50\%}+1}}$<br>Spearman-Kärber:<br>$\log_{10}(\hat{N}_0) = d_{100\%} - \left[ \frac{1}{2} - \sum_{d=d_{100\%}}^{d_{\max}} \frac{E_d}{T} \right] \log_{10} D$ | $0 = \sum_{d=1}^{d_{\max}} \frac{E_d - T + T \exp\left(-\frac{\hat{N}_0}{QD^d}\right)}{QD^d \left(1 - \exp\left(-\frac{\hat{N}_0}{QD^d}\right)\right)}$<br>Initial guess:<br>$\hat{N}_0^{\text{init}} = \frac{-QD^{d_c}}{2} \ln \left[ \left(1 - \frac{E_{d_c}}{T}\right) \left(1 - \frac{E_{d_c+1}}{T}\right)^D \right]$ |
| Luciferase Reporter ($\mu = \frac{N}{M}$) | $\hat{\mu} = \frac{1}{L_0 MT} \sum_{t=0}^T L_t^{\text{data}}$ | $0 = \frac{1}{T} \sum_{t=0}^T L_t^{\text{data}} - L_0 f_{\max} M e^{-\hat{\mu}} \sum_{r=0}^{N_0} \frac{r \hat{\mu}^r}{(K+r)r!}$<br>Initial guess:<br>$\hat{\mu}^{\text{init}} = \frac{1}{L_0 MT} \sum_{t=0}^T L_t^{\text{data}}$ |

VQAs are primarily concerned with inferring *N*_0_ and assume *a priori* knowledge of *M* and the particle to PFU ratio *Q*. In actuality, there can be variability in the number of cells present in the microtiter well and, as discussed in Section IIA, the true value of *Q* is dependent on the particular protocol and particular conditions under which an assay was performed. If an alternative assay (RNA tagging, spectroscopy, super-resolution imaging, etc.) not using cell infection can accurately measure *N*_0_, then, in theory, a subsequent infection assay can be used to infer a more reliable measure of *Q*. In fact, in our analysis of the plaque assay presented in Appendix E of the SI, we show that one can determine a significantly higher amount of information about *Q* with the same assay protocol if *N*_0_ is *a priori* known, rather than the reverse case. Thus, one may argue that assays that employ serial dilution, such as plaque and endpoint dilution assays, may be better utilized to infer *Q*. Because the underlying likelihood of the data in all assays would be the same, the same derivation techniques would follow with respect to *Q* in order to formulate its maximum likelihood estimate. This analysis shows the robust utility of a full probabilistic model and data likelihood function.

Although the derived assay models provide explicit equations for inference, many of the expressions are analytically unsolvable and require numerical solutions. To improve the accessibility of some of our results, we have created a web-based tool (available at https://bamistry.github.io/SMOI/) that can accept data from either plaque, endpoint dilution, or luciferase reporter assays and automatically estimate the parameter of interest. Ultimately, these tools can be used for analysis of future virological studies, but may also be useful when revisiting results of studies that stress quantifying viral infectivity [15, 41]. For studies that use serial dilution assays, our approach stresses the advantages of using information in the data associated with *all* dilution numbers rather than just that of the critical dilution.

Our probabilistic models of viral infection can be further generalized to include, for example, the effects of cell size inhomogeneity, coinfection, and viral interference. In the Supplemental Information, we provide a framework that would allow one to explore how these confounding factors can further alter the signal of a virus assay. Future refinement of these extensions can help to ultimately derive a mechanistic model for the probability of a single virus successfully infecting a host cell, which we defined as *Q*^−1^. Understanding this probability of infection can help aid further experimental design and allow better quantification and resolution of the infection dynamics of particular viral strains.

## Author Contributions

BM derived mathematical formulae, developed statistical inference framework, performed simulations, generated plots, and wrote the initial draft. MRD and TC verified the mathematical results, contributed to their analyses, and edited the manuscript. TC conceptualized, designed, and supervised the research.

## Acknowledgments

This work was supported in part by grants from the NSF (DMS-1516675) and the Army Research Office (W911NF 14-1-0472). We are especially grateful to Dr. Nicholas Webb, Prof. Benhur Lee, and Prof. Jerome Zack for insightful discussions.

## SUPPLEMENTARY INFORMATION

## APPENDIX A: MATHEMATICAL APPENDICES

### SMOI Probability

To derive Eq. 2, we index all cells with 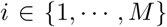 and define 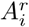 as the event that cell *i* is infected by exactly *r* IUs. Then, given *N* IUs across all *M* cells, the probability of 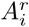 is given by

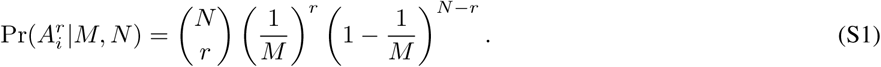

Since cell sizes are assumed to be homogeneous, the probability in Eq. S1 is the same for all cells, but the events 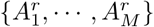 are not independent as the number of IUs *N* shared among the *M* cells is finite. Thus, we use the inclusion-exclusion principle [40] to derive

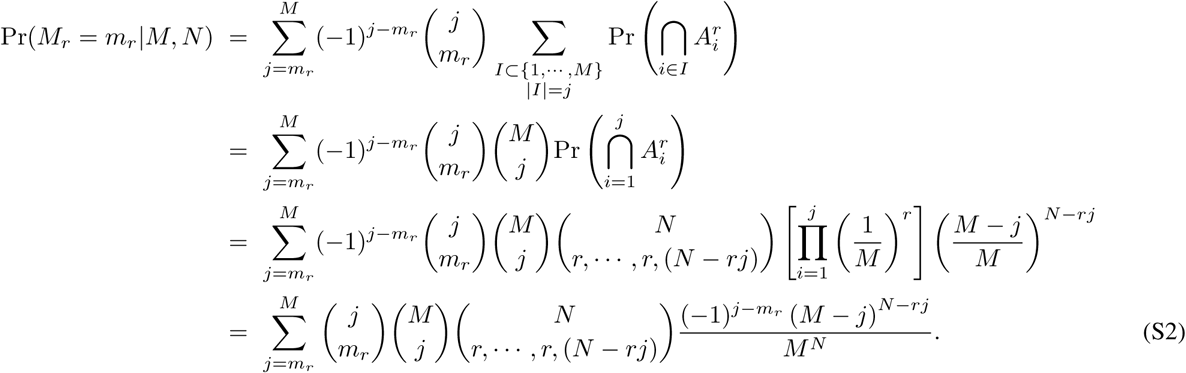

Note that the inner summation in the first identity above is over every possible collection of cells of size *j*, but as each cell is identical, the sum can be reduced to a single joint probability with the binomial degeneracy 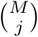.

### Expected Value and Variance

For the generalized *c*-th moment 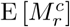 of the number of cells *M_r_* infected by exactly *r* viruses, we start with Eq. 2 to obtain

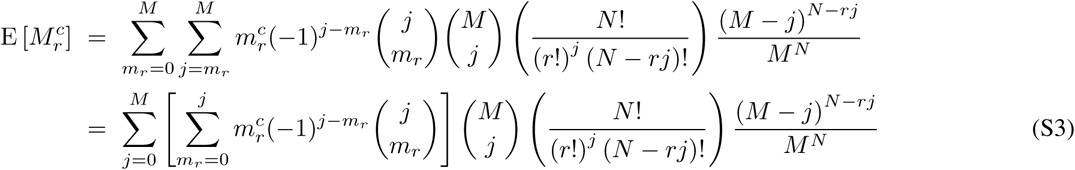

To aid our derivation, we define the function *u*(*j*,*c*) as

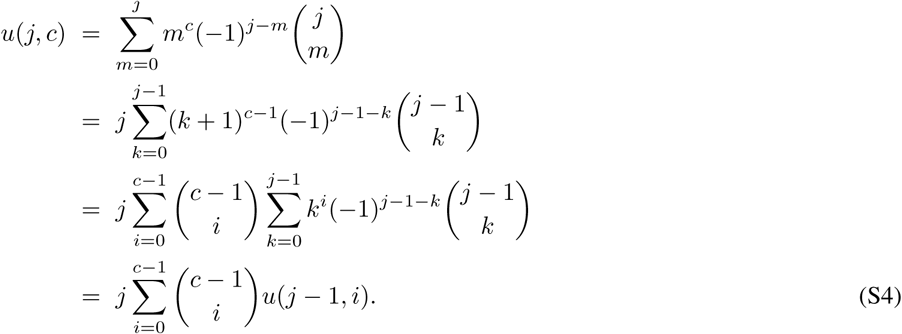

This is a recursive relationship from which we can evaluate any *u*(*j*,*c*) using all *u*(*j* – 1, *i*) such that 0 ≤ *i* < *c*. We evaluate the first three cases *u*(*j*, 0) = *δ*_0,*j*_, *u*(*j*, 1) = *δ*_1,*j*_, and *u*(*j*, 2) = *δ*_1,*j*_ + 2*_δ_*_2,*j*_, where *δ*_0,*j*_ is the Kronecker delta operator that returns the value 1 when the two subscript arguments are equal and 0 otherwise. We use the result for *c* = 1 and Eq. S3 to calculate the expected value of *M_r_* as

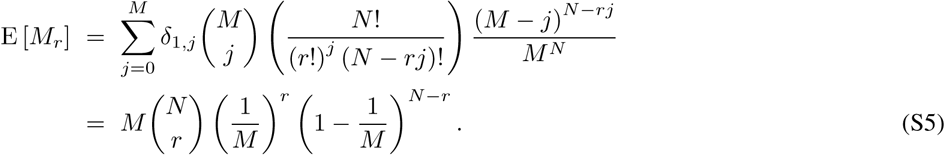

We obtain the second moment 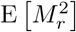 using the same method in order to obtain the variance of *M_r_* as

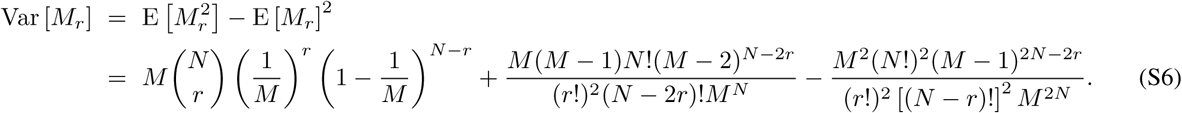

### Asymptotic Approximation

For the derivation of Eq. 6, we take the mathematical limit *N*, *M* → ∞ while keeping the ratio 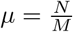 fixed and approximate Eq. 2 as follows:

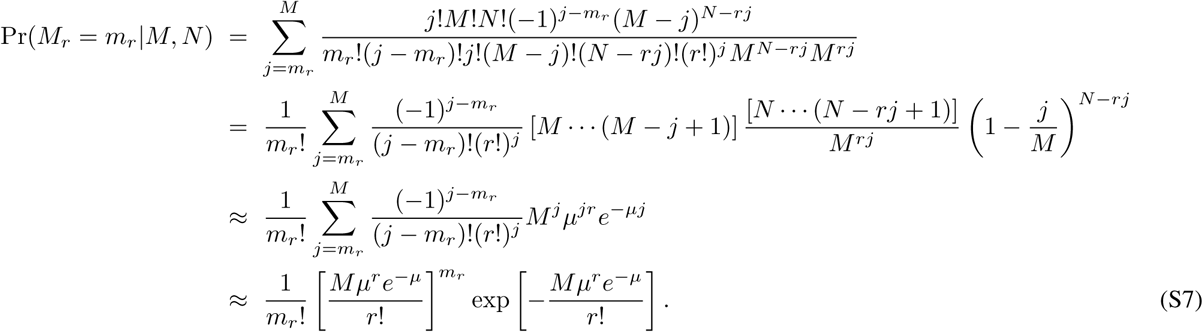

Note that, although the first approximation requires *j* in the summation to be sufficiently smaller than *M*, any contribution from the summation for *j* close to *M* vanishes due to both the (*j* – *m_r_*)! term in the denominator and the 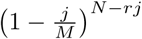 term approaching 0. Under the same large *M*, *N* limit, we can derive an asymptotic approximation of the joint probability distribution by taking the natural log of both sides of Eq. 5:

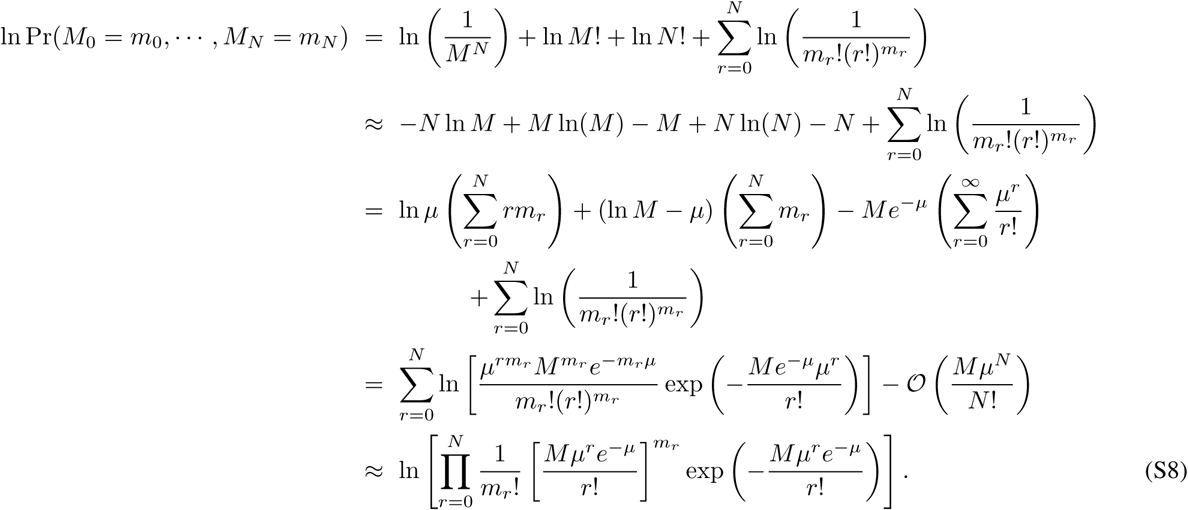

Since the argument in the right-hand-side of the last approximation is the same as Eq. 6, we arrive at the result in Eq. 8.

### Number of Infected Cells

To derive Eq. 9, we first define *N_d_* as the number of virus particles present in the viral solution after dilution of a factor of *D^d^*. Obtaining *N_d_* is effectively analogous to taking a volume of the initial viral stock scaled by *D*^−*d*^ and counting the number of particles captured in the volume. Thus, we expect *N_d_* to be Poisson-distributed with mean *N*_0_*D*^−*d*^ and discrete probability density function given by

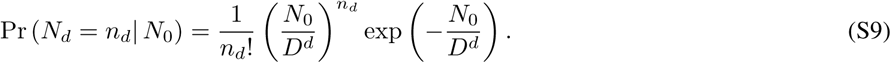

Once *N_d_* is chosen from the above distribution, for a given “particle to PFU ratio” *Q*, the number of IUs *N* follows a binomial distribution with a probability function similar to Eq. 1, but with *N*_0_ replaced with *N_d_*. Note that, given an SMOI 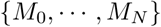, it is immediate that *M\** = *M* − *M*_0_. Using this modified density of *N* and Eqs. 2 and S9, we can derive the discrete probability density function of *M*\* at a given dilution number *d* as

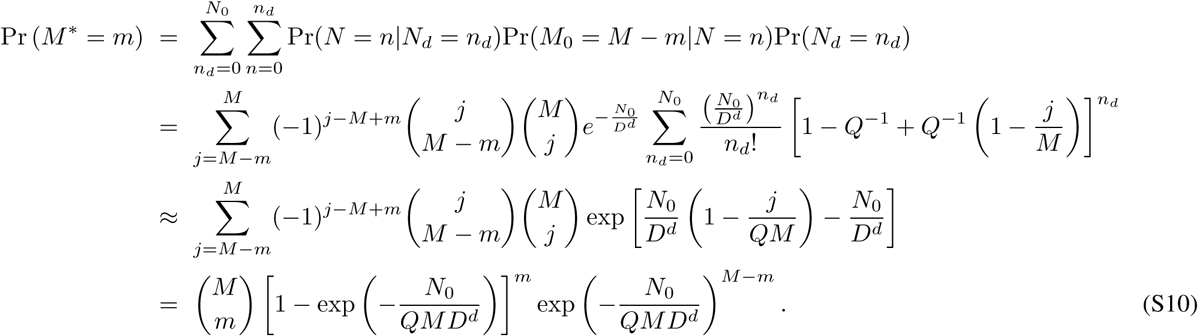

Note that the approximation that closes the exponential term in the final result employs the assumption that *N*_0_ is sufficiently large.

## APPENDIX B: INHOMOGENEOUS CELL SIZE

We derived the probability distribution in Eq. 2 assuming the plated host cells are of identical size and volume. This may not necessarily be the case as each cell exists at different stages of the mitotic cycle, will attach to the plate bottom at random locations, and contain deformities in shape and size. Assuming cells cover the entire surface of the well bottom, Pineda et al. [42] showed that the cell size proportion *p_i_* for cell *i* is gamma distributed with probability density

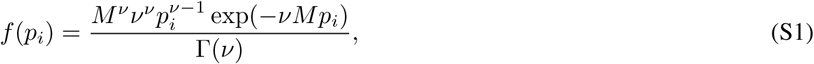

where *ν* is a parameter that can be estimated, for example, by fitting imaging data of cells. Under a specific realization of cell size distributions 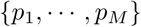, we define 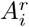 as the event that cell *i* is infected by exactly *r* viruses with probability

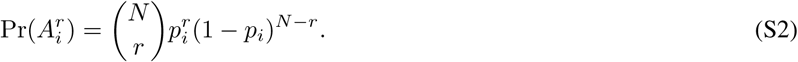

Using the inclusion-exclusion principle as above, we derive the conditional probability distribution of the number of cells *M_r_* that were infected by exactly *r* viruses as

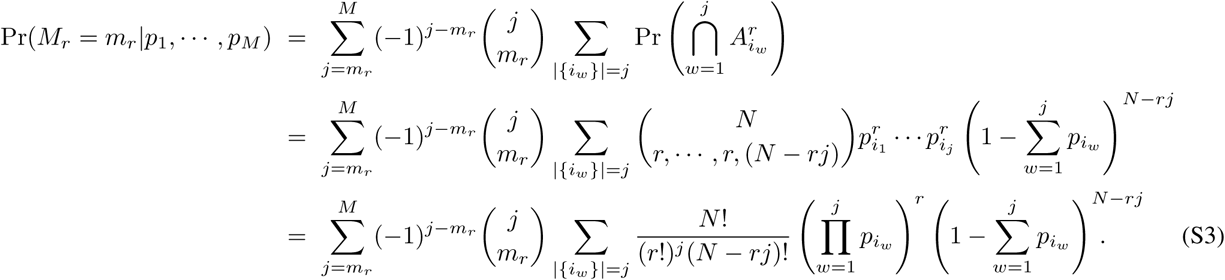

In order to obtain the full probability, we first take note that each cell size proportion *p_i_* is dependent on each other as they are constrained by 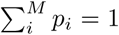. We avoid this dependency by noticing the expression in Eq. S1 approaches zero very rapidly as *p_i_* moves away from the expected value 1*/M*. If we define a sufficiently large proportion 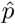 such that the interval 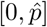 contains the majority of the area under the probability density in Eq. S1, we can make the approximation

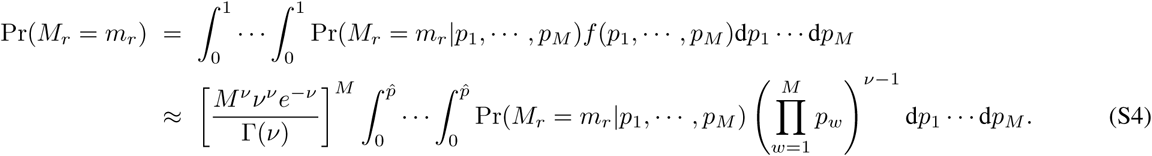

It is clear that introducing cell size inhomogeneity dramatically increases the complexity of our probabilistic SMOI model. For relatively small numbers of cells *M*, image processing can be used to determine an estimation of a particular realization of cell size distribution 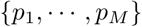 for a given experiment and factored into Eq. S3. Note that once the probability distribution of cell counts 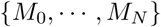 is determined for a given realization of cell sizes 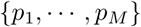, all subsequent analysis and derivations follow the same way as in the homogeneous cell size assumption.

## APPENDIX C: COINFECTION

As a vector for infection, the primary function of a single virus particle is to deliver its genetic contents into the host cell cytoplasm or nucleus [10–12]. The typical model for viral infection assumes each virus contains all the genetic material required to replicate within a host cell [14, 15]. Certain plant and fungi viruses, however, require two or more particles to successfully replicate within a host cell since each particle contains only part of the complete genome [43]. Similarly, RNA viruses that target animal cells undergo error prone replication, resulting in partially complete genome sequences. These damaged viral genes may encode proteins needed for the host cell to successfully replicate new viruses. In this case, regardless of a successful viral infection, new viruses capable of infecting further host cells will not be produced. Additional viral infections that contain the missing sequence fragments, though, can “rescue” the cell’s ability to replicate the virus, a phenomenon known as coinfection. In the context of our definition of SMOI, we now make the distinction between *M_r_*, the number of cells that have been infected by viral genomes from exactly *r* distinct virus particles, and 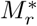, the number of cells that are fully capable of replicating new functioning viruses upon undergoing *r* distinct viral infections. It is immediate that each 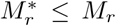 and their sum 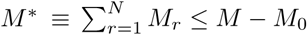, so the results in Eqs. 9 and 12 are not sufficient to quantify the total number of virus-producing cells.

In order to model coinfection, we need to consider the genome of the virus species of interest. Specifically, we assume the genome is made up of *G* distinct genes. For example, many variants of HIV-1 carry a gene sequence containing *G* = 9 genes [10]. In our model, we assume each gene encodes a protein that is essential for replication. Though individual nucleotide changes due to random mutations may result in an amino acid chain that is no longer functioning, some genes may be robust to these changes due to codon degeneracy or the gene’s shear length [44]. Thus, we assume each gene 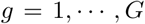 contained within a viral particle has a probability *q_g_* of losing function. If a cell is infected by exactly *r* viral genomes, we define 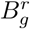 as the event that gene *g* is still no longer functional, so that 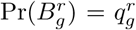. To quantify the probability that *k* genes are no longer functional in a host cell that has been infected by exactly *r* viral genomes, we use the inclusion-exclusion principle [40] to derive

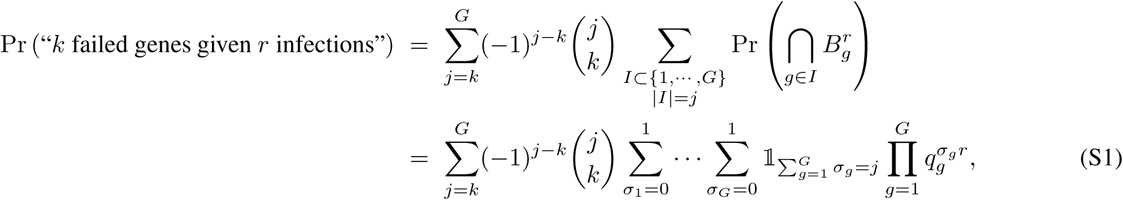

where 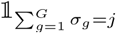 is anindicator function that returns zero when the number of nonzero *σ_g_* is not exactly *j*. The infected cell is only capable of producing viable viruses if none of the genes have failed and is equivalent to setting *k* = 0 in Eq. S1. Then we define the probability *H_r_* that a cell infected by exactly *r* viral genomes will successfully produce new viruses as

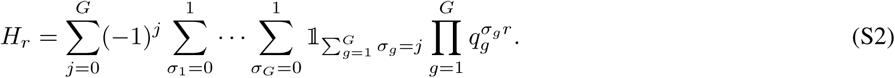

Note that the probability that a cell not infected by any viral genome will produce viruses is *H*_0_ = 0. Then, given an SMOI 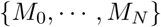, the number of cells 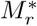 capable of virus replication after being infected by exactly *r* viral genomes is binomially distributed with parameters *M_r_* and *H_r_*. The probability of *M\** cells producing viruses is given by

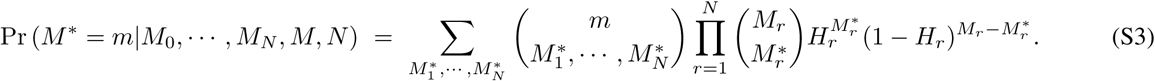

If we let *m* = 0 and sum over the density in Eq. 5 for all possible SMOI, given an IU count *N*, we can derive the probability of observing a cytopathic effect as

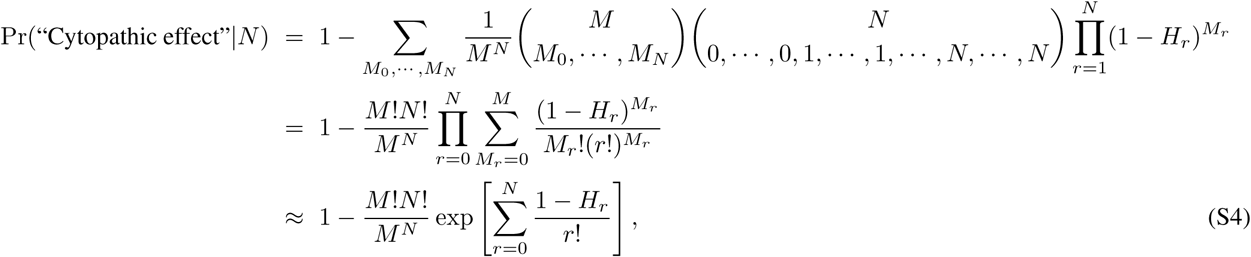

where the approximation is due to the assumption that the number of cells *M* is large. For intermediate values of *N*, computing the summation in the exponential is numerically viable, assuming the probabilities of gene failure 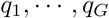 are known. Though this expression may be used in place of Eq. 12 to analyze some virus quantification assays, for large values of *N*, numerically evaluating *H_r_* becomes computationally expensive.

## APPENDIX D: VIRAL INTERFERENCE

To infect healthy cells, all species of viruses must undergo a series of events including cell attachment, entry via membrane fusion or endocytosis, and intracellular transport. Retroviruses, such as HIV-1, must also undergo reverse transcription, nuclear pore transport, and DNA integration in order to use the host cell’s transcription machinery to produce viral protein. In the models developed in this paper, the probabilities of success for each of these processes was assumed to be subsumed into the *a priori* estimated particle to PFU ratio *Q*. However, for certain retroviruses, it has been observed that after an initial infection, subsequent infections from the same virus species become less likely [45, 46]. This phenomenon, known as viral interference, is often due to the host producing new viral proteins after a refractory period that can inhibit one or more of the intracellular processes leading to integration of subsequent viral infections. To include this dynamic into our models, we first decouple the probabilities of integration from *Q* and define *N* as the number of viruses that have successfully completed viral entry into the host cytoplasm, but before all intracellular processes that lead to integration. Note that all of our results concerning the statistical multiplicity of infection (SMOI) still hold and we make the distinction between the number *M_r_* of cells infected by *r* of the *N* infectious units and the number 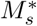 of cells with exactly *s* integrations. Furthermore, some species of virus can contain multiple copies of their genome, such as HIV-1 which contains two copies per particle [10]. Let *a* be the number of genomes contained in a single virus particle to be integrated into the host cell. Then the maximum number of possible integrations for a cell from *M_r_* is *ra*. Let *p_s_* be the probability of a viral genome integrating into the host DNA given that *s −* 1 integrations have already occurred. Define *H_r_*_,*s*_ as the probability a cell contains *s* successful integrations given that it was infected by exactly *r* distinct virus particles and is given by

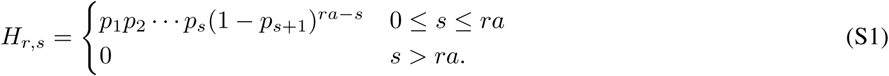

If we define 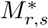 as the number of cells with *s* integrations after infection by exactly *r* virus particles, then given an SMOI 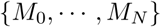 and *N*, we can derive the probability function

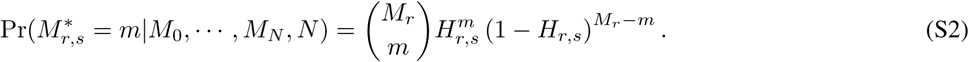

Noting that 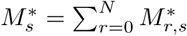 is the number of cells with exactly *s* integrations, we can use Eqs. 6 and S2 to derive the expected value as

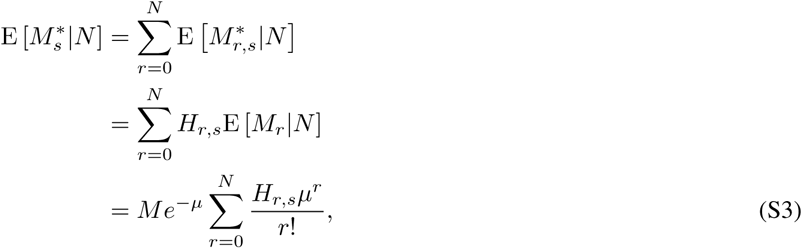

where 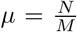. Note that if we are concerned with the total number 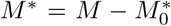 of cells with at least one integration, as is the case for the probability distributions derived for assays employing serial dilution, issue of viral interference is negligible, allowing us to subsume the probability of the first integration into the particle to PFU ratio *Q* as before and leave all subsequent virus quantification analysis unchanged from the results in Section III A and III B. However, for assays that attempts to quantify the total number of integrations, such as the luciferase reporter assay, the expectation in Eq. S3 can be used, assuming the probabilities 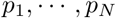 have *a priori* been estimated.

## APPENDIX E: SENSITIVITY ANALYSIS

The probability models derived in Section II A allowed us to construct the likelihood functions for the plaque, endpoint dilution, and luciferase reporter assays in Eqs. 14, 20, and 27 for the primary purpose of inferring unknown parameters such as *N*_0_ and *µ*. The utility of these functions can be extended to performing sensitivity analysis on these maximum likelihood estimates (MLE) and optimizing experimental design. This requires constructing a Fisher Information Matrix (FIM), a quantitative measure of the information one can extract for a likelihood function with an arbitrary set of data [47, 48]. The FIM, which we will denote as *J*, is constructed by computing the gradient of the log of the likelihood function with respect to the parameters being inferred. For example, for the plaque assay and potentially inferred parameters *N*_0_, *Q*, and *M*, *J* is given by

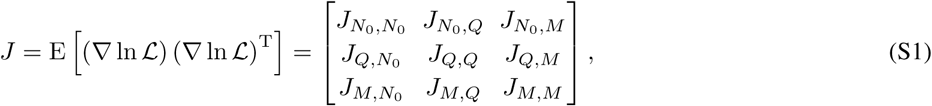

where we derive

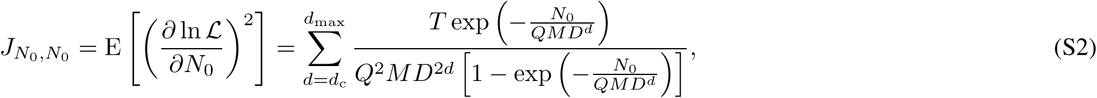

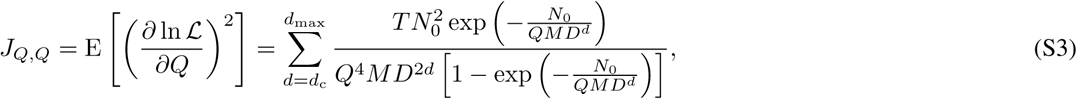

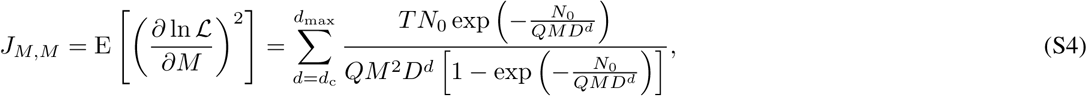

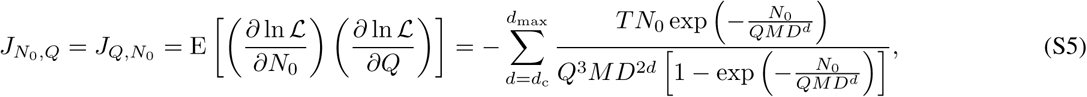

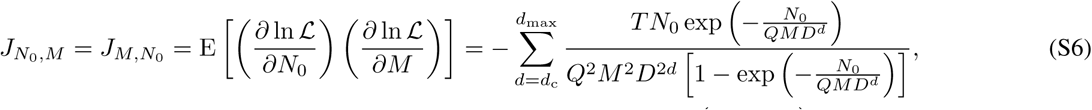

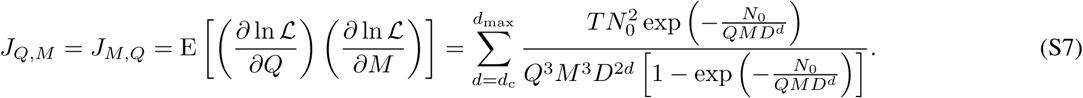

In particular, the elements of the main diagonal of *J*, known as Fisher Information Numbers, are interpreted as the “precision” of each MLE and can inform an experimentalist of the potential variation in their inferred parameter with respect to data defined by the likelihood function. Comparing the main diagonal elements can offer insight into experimental design. To illustrate, in the example above, it is immediately apparent that the ratio of *J_Q_*_,*Q*_ to 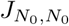 is 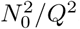, where it is understood that *N*_0_ is typically several orders of magnitude higher than *Q*. This implies that the likelihood function of Eq. 14, and, by extension, the plaque assay itself contains far more information about the parameter *Q* than *N*_0_. This provides an analytical way to decide which parameter estimation should be the focus of a particular assay.

A more general use for the FIM is to understand the variance of an MLE given an arbitrary set of data. Independent, but identical experiments can produce different estimates for each parameter and, according to the Cramer-Rao inequality, the matrix inverse *J*^−1^ will provide a theoretical lower bound on the covariance matrix of the parameter estimates [47]. Furthermore, it can be shown that the distribution of MLEs asymptotically approaches a normal distribution centered around the true experimental parameter value with covariance *J*^−1^ as the amount of data increases [49]. For single point estimation, the FIM reduces to the one Fisher Information Number with which the reciprocal can be used to approximate the variance of a parameter. For example, the plaque assay is typically used to infer only the parameter *N*_0_, so using Eq. S2, we can obtain the asymptotic approximation

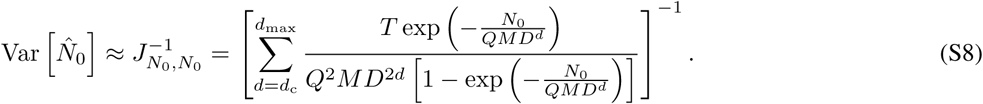

This analytical expression for the variance can be used to determine confidence intervals of the MLE, perform sensitivity analysis of other parameters, and aid in optimal experimental design.

